# Inhibition of *Streptococcus pneumoniae* autolysins highlight distinct differences between chemical and genetic inactivation

**DOI:** 10.1101/2020.09.16.300541

**Authors:** Brad A Haubrich, Saman Nayyab, Caroline Williams, Andrew Whitman, Tahl Zimmerman, Qiong Li, Yuxing Chen, Cong-Zhao Zhou, Amit Basu, Christopher W Reid

## Abstract

Despite renewed interest, development of chemical biology methods to study peptidoglycan metabolism has lagged in comparison to the glycobiology field in general. To address this, a panel of diamides were screened against the Gram-positive pathogen *Streptococcus pneumoniae* to identify inhibitors of bacterial growth. The screen identified the diamide **fgkc** as a narrow spectrum bacteriostatic inhibitor of *S. pneumoniae* growth with an MIC of 7.8 μM. The diamide inhibited detergent-induced autolysis in a concentration dependent manner indicating peptidoglycan degradation as the mode-of-action. Genetic screening of autolysin mutants suggested LytB, an endo-N-acetylglucosaminidase, involved in cell division as the potential target. Surprisingly, biochemical, and phenotypic analysis contradicted the genetic screen results. Phenotypic studies with the *Δlytb* strain illustrate the difference between genetic and chemical inactivation of autolysins. These findings suggest that meta-phenotypes including autolytic activity, cell morphology, and genetic screening can be the result of the complex interaction of one or more possible pathways that are connected to cell wall metabolism.

## Introduction

Antibiotic resistance is a growing global threat. Drug-resistant *Streptococcus pneumoniae* alone is estimated to cause 1.2 million infections with an excess of 96 million USD in medical costs per annum.(1) In light of this, there is need for the development of new therapeutics. Peptidoglycan (PG) is the primary structural heteropolymer conferring strength and cell shape determination in both Gram-negative and Gram-positive organisms (Figure 1). The PG backbone is composed of β-1,4-linked N-acetylmuramic acid (MurNAc) and N-acetylglucosamine (GlcNAc). Attached to the C-3 lactyl moiety of MurNAc is a stem peptide that is involved in cross-linking the adjacent glycan strands to form the three-dimensional structure. The incorporation of new material into the existing cell wall requires the delicate homeostasis of biosynthetic and degradative enzymes to bring new material into the stress bearing layer to prevent lysis.(2, 3) Disruption of this interplay between degradative and biosynthetic processes via chemical inhibition could provide unique insights into their biological role. Deciphering the exact biological role of autolysins has been a formidable task as functional redundancy complicates attribution of biological roles. Recent biophysical(3, 4) and computational studies(5) of bacterial autolysins have begun to unravel their roles in the release of stress in the cell wall to allow for incorporation of new material. A renaissance in peptidoglycan metabolism research has started to provided new chemical biology tools to study synthesis (6) and further insight into membrane mediated steps (7–9) and the role endopeptidases play in methicillin resistance.(10) While the cell wall and peptidoglycan in particular have provided a wealth of clinically relevant antimicrobial targets (11), our understanding of the complex interplay between degradative and synthetic steps is still developing.

**Figure 1.**
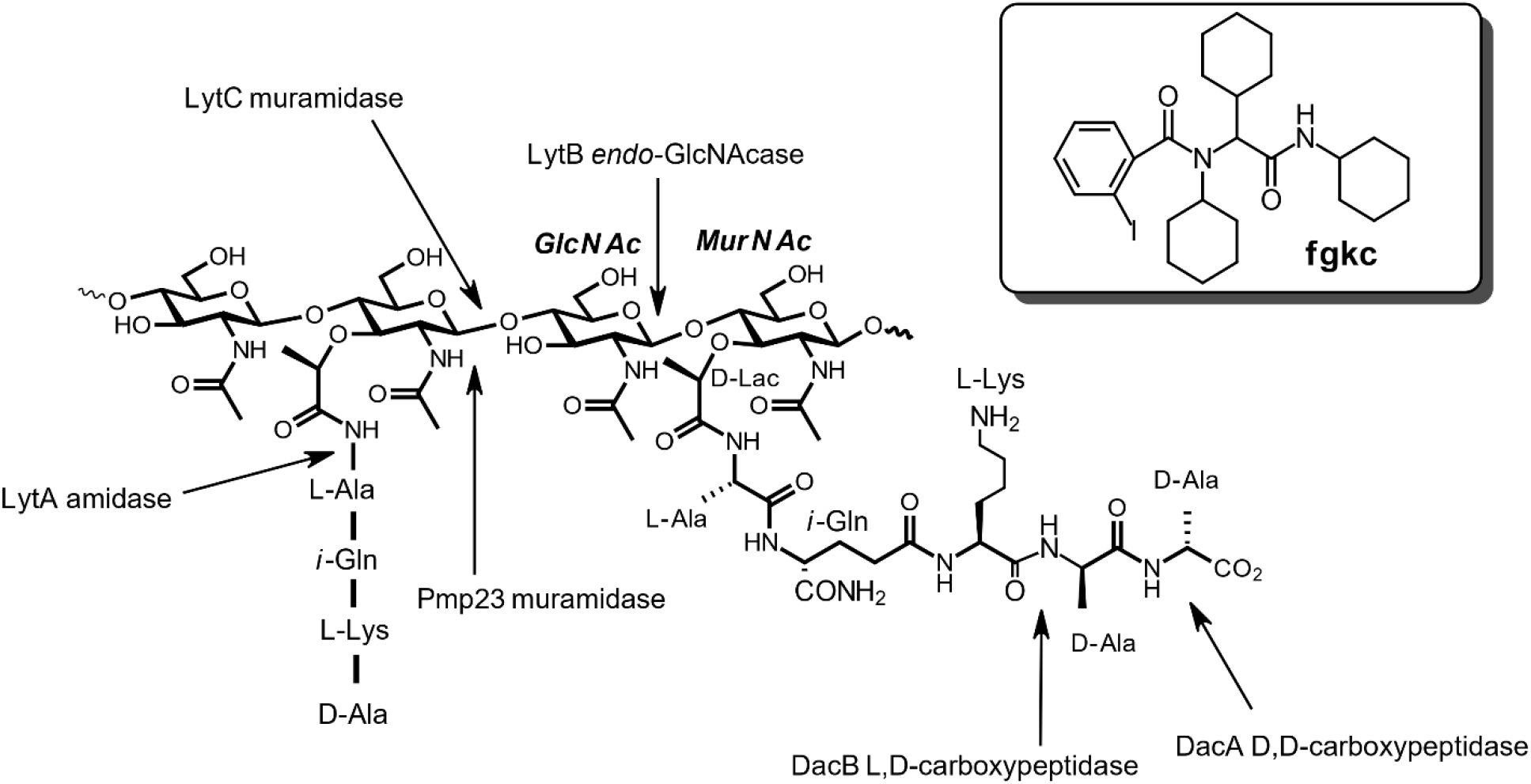
Structure of peptidoglycan showing the cleavage sites of several of the characterized autolysins in *S. pneumoniae*. Inset: structure of the antimicrobial diamide **fgkc**.

Previously, we had carried out a screen of 21 diamides for anti-bacterial activity against the Gram-positive *Bacillus subtilis*. This screen identified the diamide **fgkc** (Figure 1 inset) as a bacteriostatic inhibitor of *Bacillus subtilis* growth that targets the major active *N*-acetylglucosaminidase (GlcNAcase) LytG *in vitro*.(12) Here we report on the screening of this panel of diamides against *S. pneumoniae,* identifying **fgkc** as a bacteriostatic inhibitor of cell growth. A series of subsequent mode-of-action studies in *S. pneumoniae* highlights distinct differences between chemical and genetic inactivation of autolysins.

## Materials and Methods

### Strains and compounds

*Streptococcus pneumoniae* 6305 and R6 were purchased from ATCC (Mannassas, VA), and *S. pneumoniae* IU1945 (Δ*lytB, *Δ*lytC, *Δ*dacA,*Δ*dacB, *Δ*pmp23, *Δ*pbp1a*) mutants were kindly provided by Dr. Malcolm E. Winkler at the University of Indiana.(13) *S. pneumoniae* TIGR4 and TIGR4 *ΔlytB* strains were previously reported.(14) *S.pneumoniae* strains were grown in Mueller Hinton (MH) broth (Sigma Aldrich, St. Louis, MO) supplemented with 5 % (v/v) sheep blood (Lampire Biological, Pipersville PA) or MH agar plates containing 1.5 % (m/v) Bacto agar and 5 % (v/v) sheep blood at 37°C. *Staphylococcus aureus* was grown in MH broth or on solid media, *Clostridiodes difficile* was grown in brain heart infusion (BHI) and *Escherichia coli* DH5α on Lauria Burtani (LB) broth or solid media.

Diamide inhibitors were synthesized as described previously.(12) Other reagents, unless otherwise specified were purchased from Sigma (St. Louis, MO).

### MIC assays

MIC values were determined using the resazurin method.(15, 16) Briefly, second passage cells of *S. pnuemoniae* 6305, TIGR4, or R6 were grown in MH broth, and standardized to an OD_600nm_ = 0.4. Inhibitors were analyzed *via* serial dilution into PBS media in microtitre plates. Microtitre plates were inoculated with a 1/20 dilution of the OD_600nm_ = 0.4 cell culture with a final concentration of 1 % DMSO. Cultures were grown statically for 24 h at 37 °C, followed by addition of 30 μL of a 0.01% (m/v) solution of resazurin. The plates were incubated for 15 min to allow stabilization of color production. MICs were read directly off the plate; MICs were recorded as the lowest concentration that inhibited growth. MIC assays with *S. aureus* were performed in MH broth, *C. difficile* in BHI broth, while *E. coli* was performed in LB.

### Morphological studies of S. pnuemoniae

Cultures were prepared from second passage of *S. pneumoniae* 6305, R6 and *ΔlytB*(13) as previously described for MIC determination. Cells were chemically fixed in 20 mM HEPES pH 6.8 containing 1% formaldehyde. (17) Samples were fixed overnight at 4 °C to limit *de novo* cell wall biosynthesis during fixation. Samples were stained with 0.1% methylene blue (solution in 20% ethanol). Samples were gently heated to 60 °C for 15-20 min to bring cells to a common focal plane. Samples were visualized using bright-field microscopy with a Zeiss Primo Star microscope at 1000× magnification. Micrographs were acquired using an Axiocam ERc5s camera and Zen lite software.

### Autolysis assays

Cellular autolysis assays were performed as previously described by Cornett and Shockman.(18) Briefly, *S. pneumonia* 6305 were grown in MH broth containing 5% (v/v) defibrinated sheep blood under anaerobic conditions. Cells were harvested by centrifugation (8,000 rpm, 5 min) and washed with PBS. Cells were suspended in PBS and autolysis induced with the addition of Triton X-100 to a final concentration of 0.1% (v/v) and turbidity monitored at 600 nm over 60 min. Rates were calculated using the linear portion of the autolysis curves with the rate of autolysis in the absence of inhibitor set at 100%.

### Chain dispersing assay

Dispersion of the *ΔlytB* chain morphology with purified LytB was carried out as previously described using the TIGR4 and associated *ΔlytB* strains.(14) LytB was added to the cell suspension at a final concentration of 2μM. The final concentration of **fgkc** in the assays was 40 μM.

### DNA intercalation assays

To determine if **fgkc** is a DNA intercalator, DNA mobility shift assays were performed as previously described using BamHI-linearized pUC18 plasmid.(19) The known DNA intercalator actinomycin D was used as a control.

### DNase I assays

Degradation of the pUC18 plasmid DNA was assayed using 150 ng of linearized pUC18 plasmid.(20) Compound titrations in DMSO were added and reactions were initiated with 0.002 units of DNase I. The reactions were incubated for 15 min at 37°C before being subjected to agarose gel electrophoresis. EDTA was used as a control for DNase I inhibition.

### Lipotechoic acid detection by Western blot

Lipoteichoic acid profiles were analyzed as previously described.(21) Briefly, *S. pneumoniae* R6 cells were cultured to saturation, harvested (3000 x g), resuspended in 6 M urea, and incubated at 37oC for 5 min to solubilize proteins. Samples were standardized by total protein content and separated by SDS-PAGE (16%) and transferred to PVDF membrane. The membrane was incubated with a 1:5000 α-phosphocholine monoclonal antibody (SSI Biotech, Santa Cruz, CA). Blots were analyzed by chemiluminescence.

## Results and Discussion

We screened a previously reported (12) panel of 21 diamides against *S. pneumoniae* using the resazurin microtiter assay (Figure S1).(16) Of the 21 compounds screened, **fgkc** was identified as a single digit micromolar inhibitor of *S. pneumoniae* growth with an MIC of 7.8 μM against strains 6305, R6, and TIGR4. Analysis of the mode of growth inhibition demonstrated that **fgkc** was bacteriostatic. These results for **fgkc** are comparable to those obtained against *B. subtilis* (MIC = 3.8 μM).(12) To assess the antimicrobial spectrum of the compound, we screened **fgkc** against the Gram-positive organisms *Clostridiodes difficile, Staphylococcus aureus* and the Gram-negative *Escherichia coli.* In all cases, no antimicrobial activity was observed up to 200 μM suggesting that **fgkc** is a narrow spectrum inhibitor of Gram-positive growth. Extrapolating from *B. subtilis* activity where **fgkc** inhibits LytG, a glycosyl hydrolase family 73 (GH73) enzyme, we presumed the target was a member of GH73. To date, *S. pneumoniae* possesses one enzyme classified as GH73 (www.cazy.org) – LytB, a cell division associated endo-β-GlcNAcase belonging to cluster 4 of GH73.(14, 22–24) In contrast, *B subtilis* LytG is an exo-acting GlcNAcase active in vegetative growth belonging to GH73 cluster 2. (25)

Based on our previous findings in *B. subtilis*, we examined whether **fgkc** inhibited autolytic activity in *S. pneumoniae* (Figure 2A). Non-ionic detergent (Triton X-100) induced autolysis of *S. pneumoniae* was inhibited in a concentration induced manner by **fgkc**, suggesting that this compound does indeed target the cell wall and autolysins specifically. We followed up with a genetic screen of cell wall autolysins (Figure 1) and PG biosynthetic enzyme knockout strains to further refine the mode-of-action and identify the molecular target (Figure 2B). Conventional wisdom in this type of experiment would presume that deletion of the target should result in a reduction in sensitivity to the compound. The genetic screen demonstrated a nearly 4-fold decrease in sensitivity to **fgkc** in the *ΔlytB* mutant (13), strongly suggesting LytB was the putative target of the diamide.

**Figure 2.**
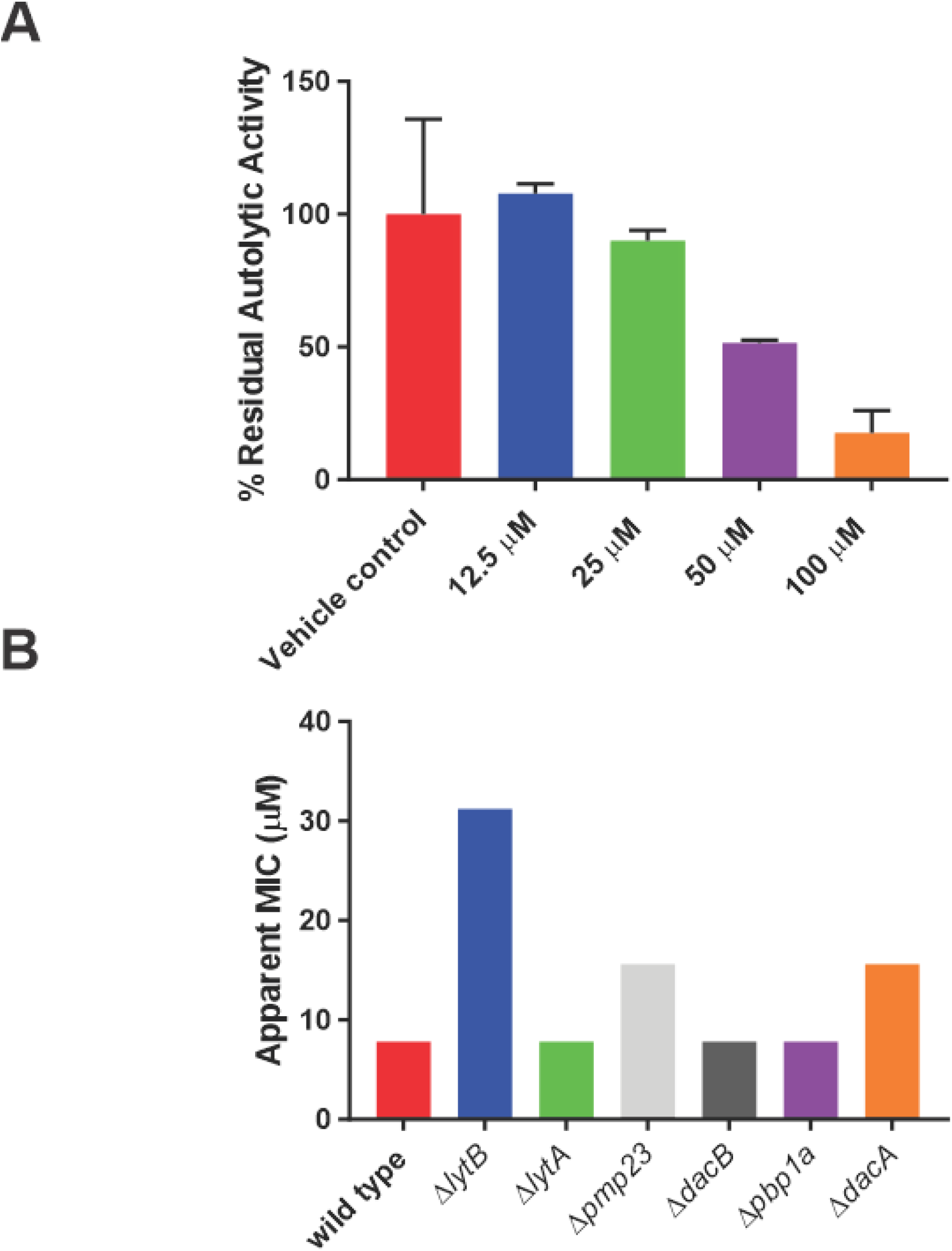
Screening of the diamide **fgkc** against *Streptococcus pneumoniae*. (**A)**The diamide **fgkc** inhibits detergent-induced autolysis. (**B**) Genetic screen of *S. pneumoniae* R6 autolysin and cell wall biosynthesis mutants (13) to identify the target of **fgkc**. Data shown is the average of experiments performed in biological and technical triplicate.

To further explore these results, morphological changes induced by sub-MIC concentrations of **fgkc** and antibiotics of known modes-of-action were investigated (Figure 3). *S. pneumoniae* treated with **fgkc** showed a change from the wild-type diploid morphology to a clumping phenotype like the cell wall acting antibiotics bacitracin and vancomycin. The morphology of sub-MIC treated cells showed a deviation from the typical coccoid shape and size. The **fgkc** induced phenotype did not correlate with the reported phenotypes of a *ΔlytB* mutant, which exhibit a chaining phenotype(14, 26, 27) A clumping phenotype was also observed with sub-MIC kanamycin. This phenotype has been associated with antibiotics that target intracellular protein synthesis.(28) Additionally, lipoteichoic acid (LTA) disruption was monitored by Western blot in the presence of sub-MIC **fgkc**. Results indicated that changes to LTA incorporation in the cell wall was not a contributor to the observed **fgkc**-induced phenotype (Figure S2). In summary, the morphological changes are consistent with the hypothesis that **fgkc** is acting on a cell wall target.

**Figure 3.**
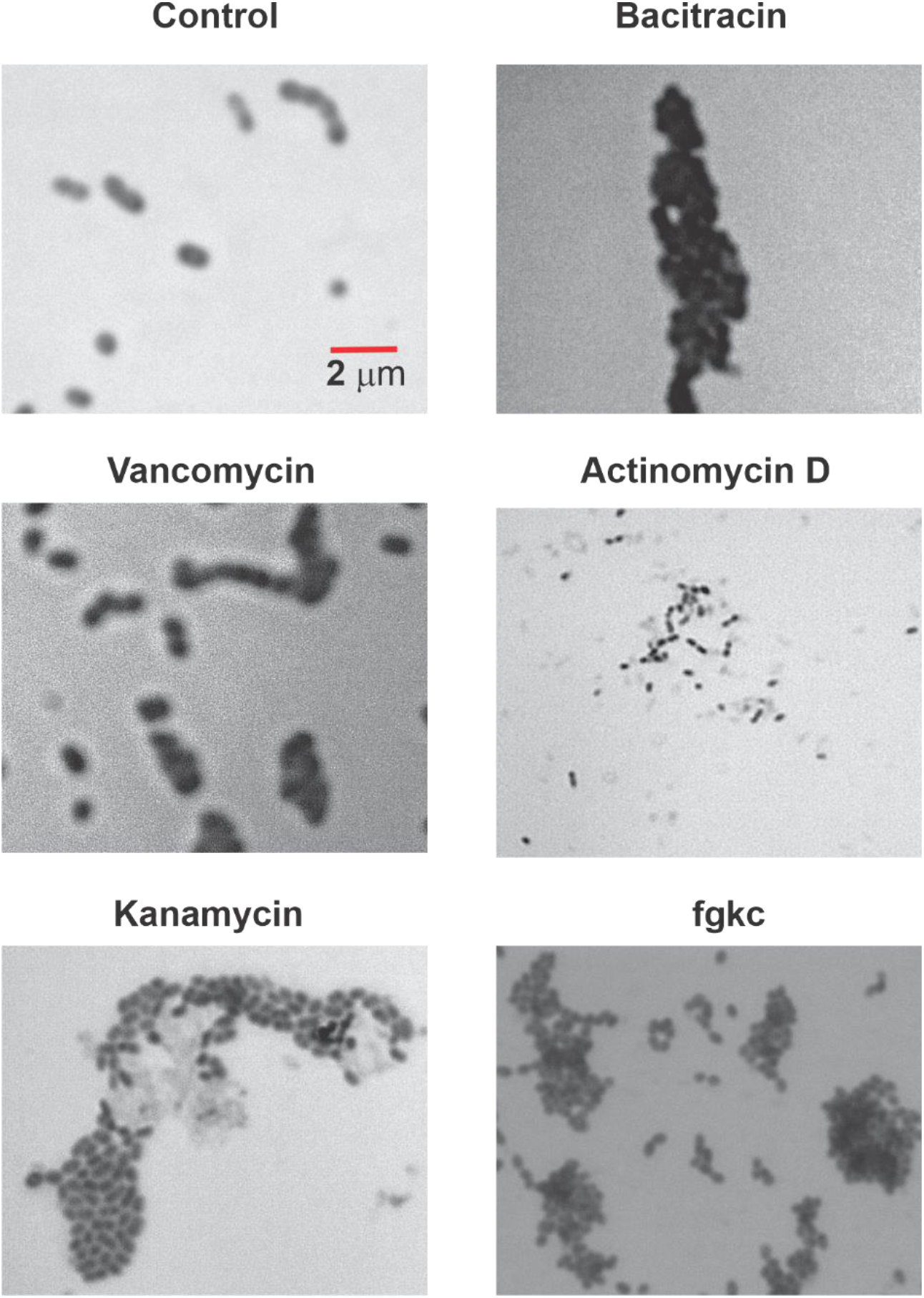
Phenotypic analysis of wild-type *S.pneumoniae* in the presence of sub-MIC (0.7x) of antibiotics with well-defined modes of action or **fgkc**. Cells were treated for 90 min with antibiotic or vehicle control, fixed in 20 mM HEPES pH 7.0, 1% formaldehyde, and stained with methylene blue.

Inhibition of LytB activity was investigated in a chain dispersing assay with the *ΔlytB* strain and exogenously added LytB.(14, 27) When the *ΔlytB* mutant is treated with exogenous LytB, dispersion of the chains is observed (Figure 4). When a 5-fold MIC (40 μM) concentration of **fgkc** was added, LytB catalyzed chain dispersion was not inhibited. In vitro analysis with Remazol brilliant blue labeled PG (14, 29) and purified LytB confirmed these results. This lack of inhibition of the biochemical activity of purified LytB is inconsistent with the larger MIC observed against the ∆*lytB* mutant.

**Figure 4.**
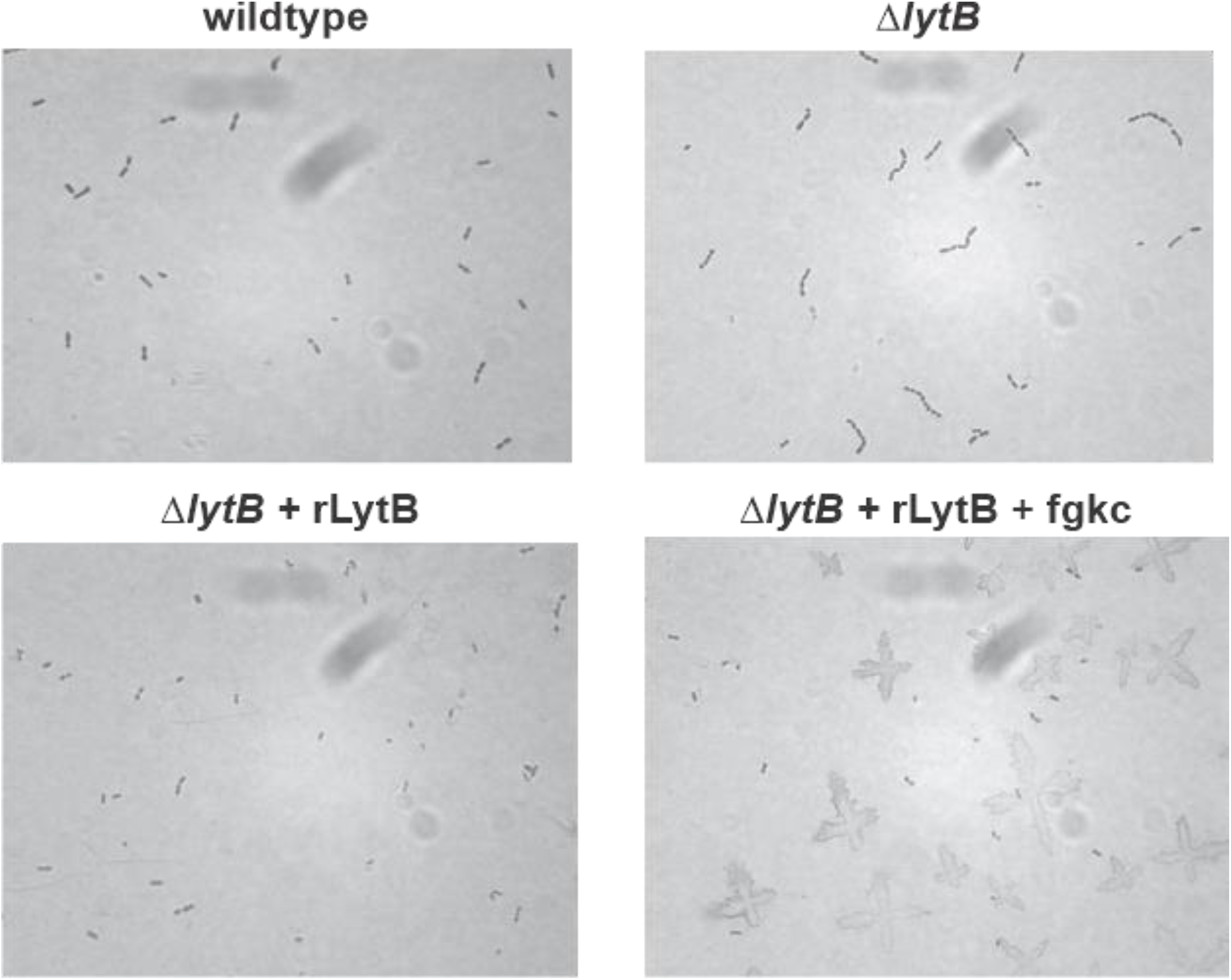
Chain dispersing assay with *S.pneumoniae *ΔlytB** strain and purified recombinant LytB (rLytB, 2 μM). In the presence of 40 μM **fgkc** dispersion of the *ΔlytB* chain phenotype is not inhibited.

This contradictory data suggests that the reduction in autolysis observed with **fgkc**, phenotypic, and genetic screening results are meta-phenotypes-a phenotype that result from the alteration of more than one pathways associated with the cell wall.(30) For instance, the clumping meta-phenotype observed in *S.pneumoniae* in the presence of **fgkc** could be generated via a direct mechanism (*e.g.* inhibition of an enzyme associated with cell wall metabolism) or an indirect indirect one (*e.g.* alterations in autolysin expression levels, changes in metabolic flux through cell wall associated pathways). To further explore this meta-phenotype hypothesis, we looked to see if the *ΔlytB*;chain phenotype could be converted to the clumping phenotype observed in wild-type *S. pneumoniae*. Sub-MIC treatment of *S.pneumoniae *ΔlytB** with **fgkc**, cefoxitin (DacA/PBP3 selective)(31, 32), or a combination of the two was examined by microscopy (Figure 5). Treatment with 0.7x MIC **fgkc** resulted in the conversion of the *ΔlytB* chaining phenotype to the clumping phenotype observed when wild-type *S. pneumoniae* is treated with **fgkc**. This observation clearly indicates that **fgkc** is perturbing the cell wall phenomena observed in the *∆lytB* phenotype, suggesting a molecular target other than LytB. *This observation also directly contradicts the genetic screen results in Figure 2B, which point to LytB as a target.*

**Figure 5.**
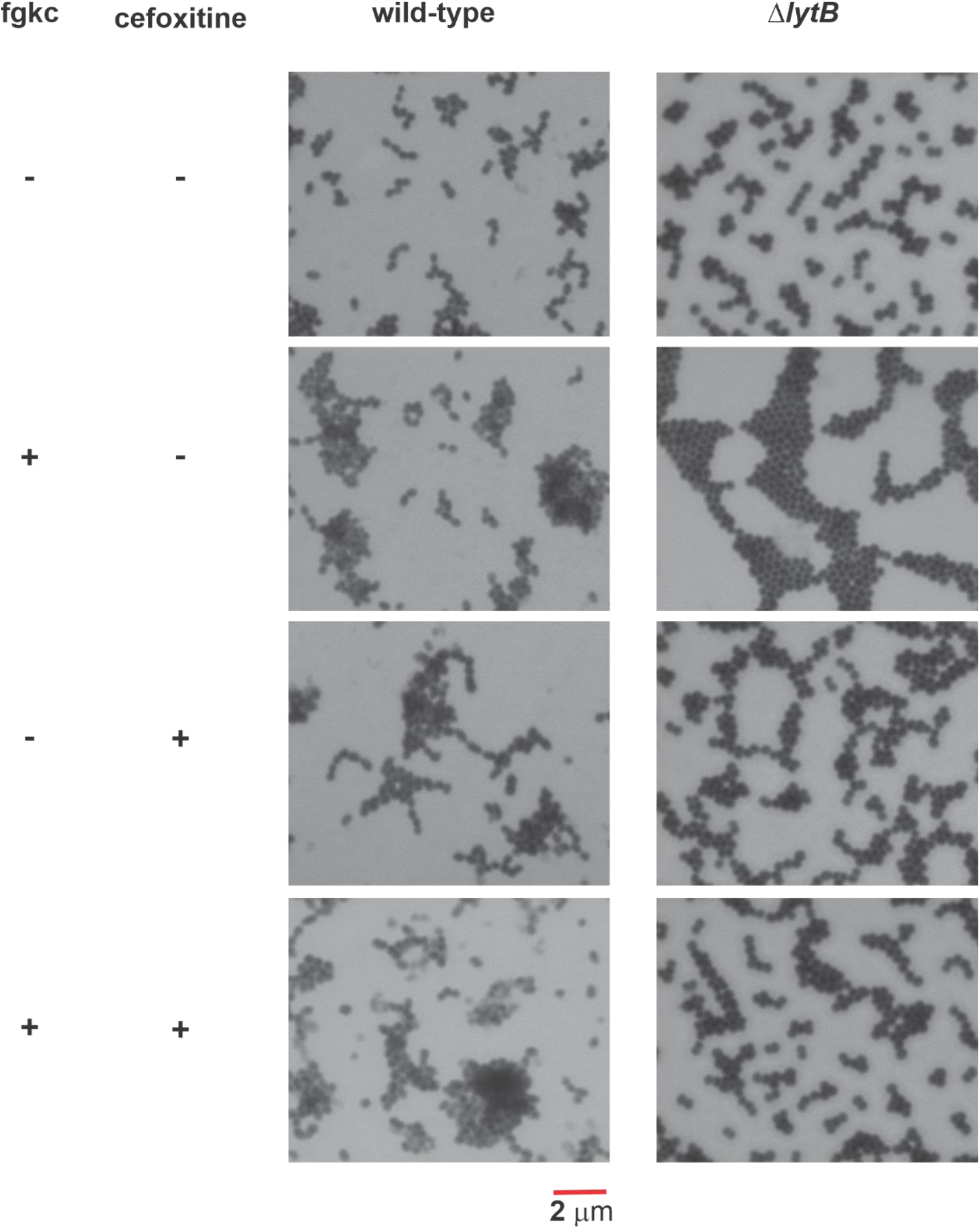
Morphological analysis of *S. pneumoniae *ΔlytB** mutant-(13) in the presence of sub-MIC **fgkc**, the β-lactam cefoxitin (DacA/PBP3 selective) or in combination. Cells were fixed in 1% formaldehyde, stained with methylene blue and visualized using bright field microscopy under oil immersion.

Interestingly, when **fgkc** was co-administered with 0.7x MIC cefoxitin a disruption of the **fgkc** – induced clumping phenotype was observed. A further analysis of the genetic screen results showed mild decreases in sensitivity (approximately 2-fold) in the *Δpmp23* and *ΔdacA* strains. Pmp23 is a putative muramidase that is required for proper localization of the Z-ring and the FtsZ-positioning protein MapZ.(33) DacA is a D,-D-carboxypeptidase (penicillin-binding protein 3) shown to be involved in cell separation.(34, 35)

Taking this data holistically, we posit that **fgkc** disrupts autolytic activity associated with the cell division machinery in *S. pneumoniae.* This hypothesis is supported by reported protein interaction data that shows the three autolysins identified in the genetic screen are associated with components of the cell division complex.(33, 36, 37) This data demonstrates that genetic knockouts of cell wall acting enzymes can be distinct from chemical knockout. This distinction has previously been observed in *Mycobacterium tuberculosis* shikimate biosynthesis.(38) Further, these data illustrate the complexity involved in deciphering the underlying mechanisms associated with these meta-phenotypes, and obfuscates target identification. Despite these challenges, the elucidation of the molecular target of **fgkc** are on-going. However, these data illustrate the promise that small molecule inhibitors to autolytic enzymes can play in furthering our understanding of peptidoglycan metabolism.

## Conclusion

The bacterial cell wall, and PG biosynthesis has provided a wealth of clinically relevant antibiotic targets. While our understanding of PG biosynthetic and cross-linking steps is fairly well established, our knowledge of the role autolytic enzymes play in the growth and maintenance of the cell wall has remained more elusive. Traditional genetic approaches to studying autolysin biological role are complicated by functional redundancy of these enzymes, where other autolysins can compensate for a loss in activity. The results presented here illustrate the complexity of PG metabolism and that phenotypes may be the manifestation of a complex interplay between associated systems. Biophysical(3, 4) and computational studies(5) have begun to elucidate the role of autolysins in relieving stress in the cell wall to allow for incorporation of new material into the stress bearing layer. The results with the diamide **fgkc** demonstrate that there are distinct differences between the genetic and chemical inactivation of cell wall acting enzymes. Collectively our data indicate that morphological, genetic, and whole cell assays (autolysis) reveal meta-phenotypes that result from the complex interaction of one or more cellular processes that are connected to cell wall metabolism. The genetic deletion of one or more autolysins disrupts the equilibrium stoichiometry of the cell wall machinery that likely results in changes to expression levels and activity to both autolytic and biosynthetic enzymes. With interest in the development of chemical biology approaches to study PG metabolism(39–43) receiving renewed attention, the diamide inhibitor **fgkc** provides an orthogonal approach to both traditional and chemical biology approaches to studying cell wall metabolism.

## Acknowledgements

Research was partially supported by the National Science Foundation (CHE2009522) and by the Rhode Island Institutional Development Award (IDeA) Network of Biomedical Research Excellence from the National Institute of General Medical Sciences of the National Institutes of Health under grant number P20GM103430 We thank J. Belval for technical assistance.

