## Supplementary Information for "Inhibition of *Streptococcus pneumoniae* autolysins highlight distinct differences between chemical and genetic inactivation"

Table of Contents

Figure S1. Structures of the diamide library.....................................................................2

Figure S2. Changes to wall teichoic acid profiles…………..………………………….……2

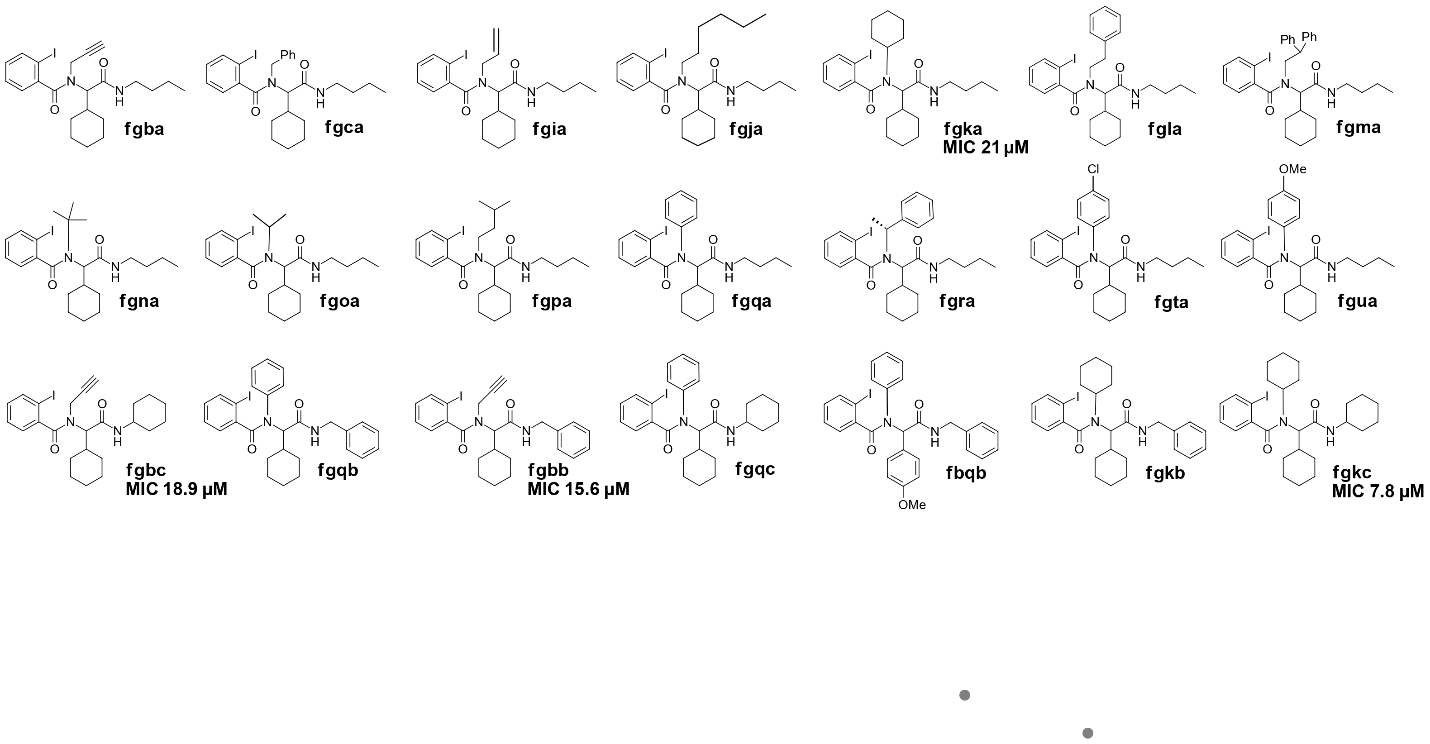
**Figure S1.** Structures of the diamide library screened against *Streptococcus pneumoniae*.

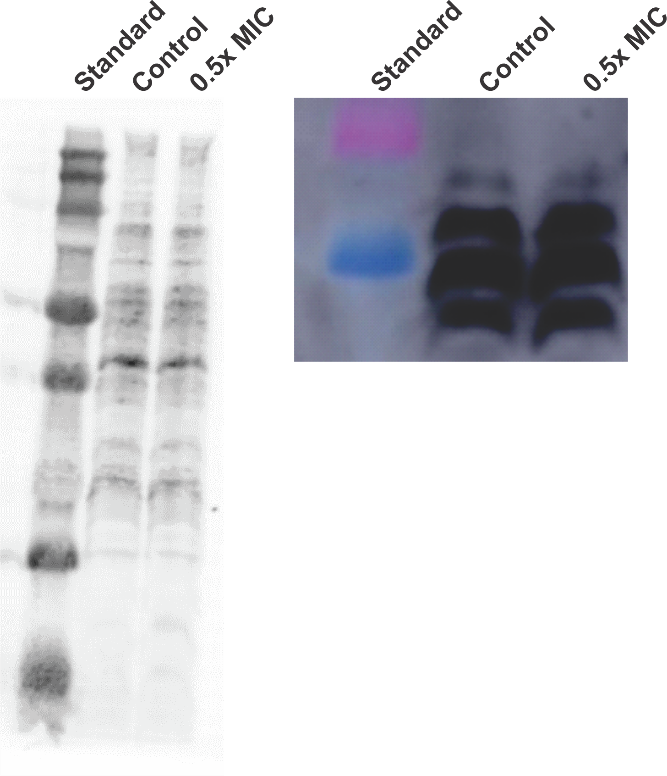

**Figure S2.** Changes in wall teichoic acid (WTA) profile (A) and changes to phosphocholine levels (B) upon treatment with 0.5x MIC **fgkc**.
